# High-content microscopy reveals a morphological signature of bortezomib resistance

**DOI:** 10.1101/2023.05.02.539137

**Authors:** M.E. Kelley, A.Y. Berman, D.R. Stirling, B.A. Cimini, Y. Han, S. Singh, A.E. Carpenter, T.M. Kapoor, G.P. Way

**Author notes:** **CORRESPONDENCE** Gregory P. Way, Tarun M. Kapoor, Anne E. Carpenter. co-senior authors (S. Singh, A.E. Carpenter, T.M. Kapoor, G.P. Way).

## Abstract

Drug resistance is a challenge in anticancer therapy, particularly with targeted therapeutics and cytotoxic compounds. In many cases, cancers can be resistant to the drug prior to exposure, i.e., possess intrinsic drug resistance. However, we lack target-independent methods to anticipate resistance in cancer cell lines or characterize intrinsic drug resistance without a priori knowledge of its cause. We hypothesized that cell morphology could provide an unbiased readout of drug sensitivity prior to treatment. We therefore isolated clonal cell lines that were either sensitive or resistant to bortezomib, a well-characterized proteasome inhibitor and anticancer drug to which many cancer cells possess intrinsic resistance. We then measured high-dimensional single-cell morphology profiles using Cell Painting, a high-content microscopy assay. Our imaging- and computation-based profiling pipeline identified morphological features typically different between resistant and sensitive clones. These features were compiled to generate a morphological signature of bortezomib resistance, which correctly predicted the bortezomib treatment response in seven of ten cell lines not included in the training dataset. This signature of resistance was specific to bortezomib over other drugs targeting the ubiquitin-proteasome system. Our results provide evidence that intrinsic morphological features of drug resistance exist and establish a framework for their identification.

## Introduction

Targeted cancer therapies are often thwarted by drug resistance (Garraway and Jänne, 2012; Pisa and Kapoor, 2020). Resistance is complex and can be categorized as acquired, manifesting in the context of prolonged treatment, or intrinsic, pre-existing in the cancer cell population (Gottesman et al., 2016). Drug resistance often results in failed therapies and cancer relapse, which makes determining the drug sensitivity of populations of cancer cells requisite for timely and effective treatment (Vasan et al., 2019). However, we currently lack unbiased methods of identifying intrinsic drug resistance in cells prior to treatment.

Bortezomib is an anticancer drug commonly used to treat multiple myeloma and nearly half of multiple myeloma patients show no initial response to bortezomib therapy, indicating intrinsic resistance (Chen et al., 2011; Gonzalez-Santamarta et al., 2020; Mitsiades et al., 2004). Malignant plasma cells in multiple myeloma depend on the timely degradation of proteins by the proteasome to prevent apoptosis (Gonzalez-Santamarta et al., 2020), making proteasome inhibitors such as bortezomib a standard of care for multiple myeloma (Hideshima et al., 2001; Vincenz et al., 2013). However, multiple myeloma is invariably fatal due to the eventual development of drug resistance (Hideshima et al., 2007).

Bortezomib resistance can be attributed to targeted mechanisms such as mutations in the bortezomib-binding pocket of the targeted proteasome subunit (PSMB5) and overexpression of proteasome subunits (Barrio et al., 2019; Franke et al., 2012; Oerlemans et al., 2008) as well as non-specific mechanisms, such as upregulation of prosurvival or anti-apoptotic pathways and enhanced cell adhesion to the extracellular matrix (Gonzalez-Santamarta et al., 2020; Hideshima et al., 2007). A priori knowledge of tumor cells’ susceptibility to candidate therapeutics could aid in identifying effective treatment options, resulting in fewer relapses and failed treatments due to intrinsic resistance. However, current methods for determining resistance status depend on viability assays, which take days to perform, or sequencing, which may be limited in its usefulness without knowledge of the full spectrum of resistance-conferring mutations (Wheler et al., 2014) or knowing specific mutations or indels in the target that suppress drug activity (Kapoor and Miller, 2017). Alternative methods for determining tumor cell susceptibility to therapy are therefore desirable.

A growing literature suggests that specific genetic alterations, treatment response, and prognosis can be predicted from conventional hematoxylin and eosin tissue slides using machine learning (Cifci et al., 2022; Lee and Jang, 2022), indicating that image data holds promise for predicting drug sensitivity. High-content screening, which uses cell-based automated microscopy to capture information-rich images, has successfully categorized small molecule inhibitors by their mechanisms and targeted pathways (Ljosa et al., 2013; Perlman et al., 2004) and shown a relationship between morphological profiles and genetic perturbations (Rohban et al., 2017), including specific mutations associated with lung cancer when in an artificial overexpression system (Caicedo et al., 2022). This screening method often uses high-throughput microscopy that generates a large amount of image data, from which thousands of quantitative, single-cell morphological features can be extracted to characterize signals that could not be discovered using low-throughput methods and would otherwise be impossible to study by eye. However, this has not been used to examine the features of drug resistance in untreated cells.

Here, we used Cell Painting (Bray et al., 2016), a multiplex, fluorescence microscopy assay that labels eight cellular components using six stains imaged in five fluorescent channels, as an unbiased method to characterize the morphological differences between untreated bortezomib-resistant and -sensitive cancer cell lines. We applied a reproducible imaging- and computation-based profiling pipeline to process the images and identify a high-dimensional cell morphology signature to predict bortezomib resistance that we evaluated using machine learning best practices. This morphological signature correctly predicted the bortezomib treatment response in seven out of ten cell lines not included in the training data and was highly specific; the signature had limited ability to identify cells resistant to other drugs targeting the ubiquitin-proteasome system (UPS) such as ixazomib, another proteasome inhibitor, or CB-5083, which targets p97 upstream of the proteasome. These results suggest that this method can specifically identify bortezomib-resistant cell lines better than random chance and establish a framework for identifying morphological signatures of drug resistance. The ability to identify drug-resistant cell lines based on intrinsic morphological features supports using microscopy to guide therapy and provides a valuable orthogonal method for characterizing drug resistance.

## RESULTS

### Isolating and capturing Cell Painting profiles for HCT116-based bortezomib-resistant clones

We first isolated and characterized drug-resistant cell lines (Fig. 1 A). We based our method on our previous work, growing a parental polyclonal cell line in the presence of the desired drug to isolate drug-resistant clones (Kasap et al., 2014; Wacker et al., 2012). To efficiently isolate drug-resistant clones, we used HCT116 parental cells that have low levels of multidrug resistance pumps and are mismatch repair deficient, providing a genetically heterogeneous polyclonal population of cells (Papadopoulos et al., 1994; Teraishi et al., 2005; Umar et al., 1994). To approximate the conditions of intrinsic resistance, we cultured these polyclonal parental HCT116 cells in a high enough concentration of a drug of interest to kill the majority of cells within days leaving a few isolated, surviving single cells in a time frame consistent with these cells harboring intrinsic resistance (Wacker et al., 2012).

**Figure 1.**
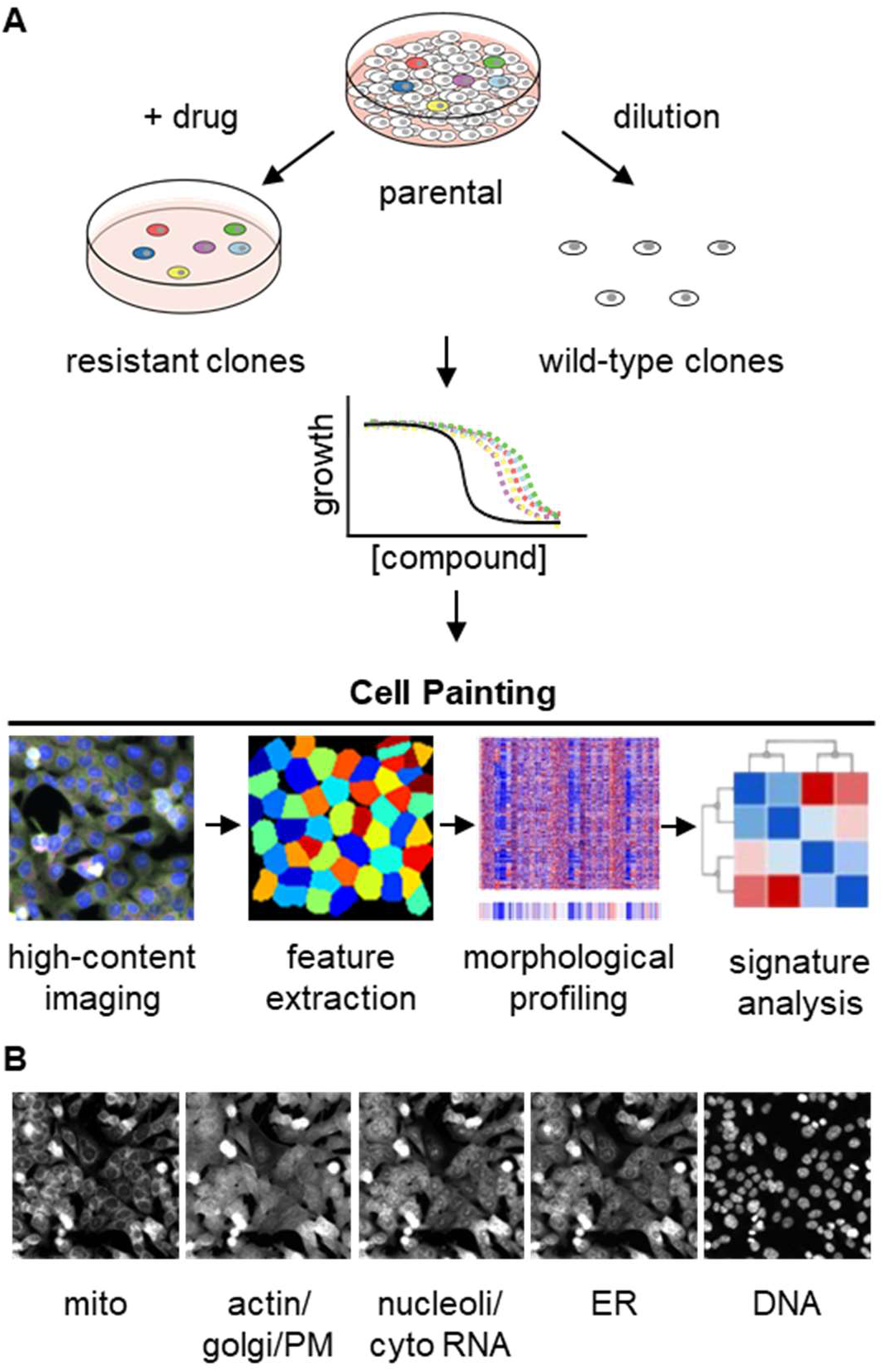
Experimental design for using Cell Painting to examine morphological profiles of drug-resistant cell lines. **(A)** Graphic of the experimental workflow for isolating and characterizing drug-resistant cell lines and then performing Cell Painting to search for morphological features of drug sensitivity. **(B)** One representative field of view of cells labeled with six fluorescent dyes and captured in five channels used for morphological profiling with Cell Painting. Each image is 230.43 × 230.43 μm.

To determine the appropriate drug concentrations to use in order to produce only a few surviving drug-resistant cells, we performed proliferation assays on HCT116 parental cells with our drugs of interest: bortezomib (proteasome inhibitor), ixazomib (proteasome inhibitor), or CB-5083 (p97 inhibitor) (Fig. 1-Supplement 1 A-D). Although our proliferation assays could not distinguish between cell death and cytostatic effects, we qualitatively confirmed cell death using brightfield microscopy. We then treated parental HCT116 cells with drug concentrations at and around the calculated LD90 for each of these small molecules to isolate drug-resistant single cells and expanded them into clonal cell lines for further experiments (Wacker et al., 2012). In addition to these drug-resistant clonal cell lines, we isolated wild-type clones by dilution of the parental line and acquired two previously isolated bortezomib-resistant cell lines (BZ clones A and E) with mutations in PSMB5 confirmed by RNA sequencing (Fig. 1-Supplement 1 E) (Wacker et al., 2012).

To screen for multidrug resistance, which might convolute any specific signature of bortezomib resistance, we measured proliferation of our clones in the presence of drugs that target disparate pathways: bortezomib (a proteasome inhibitor), taxol (a microtubule poison), and mitoxantrone (a topoisomerase inhibitor) (Liu, 1989). As multidrug resistance leads to non-specific reductions in drug sensitivity, due to drug efflux for example (Gottesman et al., 2016), we expected multidrug resistant cells to be less sensitive to bortezomib, taxol, and mitoxantrone.

When we treated our cell lines with bortezomib, only the cell lines isolated following high-dose bortezomib treatment (BZ01-10 and BZ clones A and E) had reduced bortezomib sensitivity, with LD50s ranging ∼2.8- to ∼9-fold that of the wild-type parental cell line (Fig. 1-Supplement 2 B). In contrast, the wild-type clones (WT01-05, 10, and 12-15) had LD50s ranging from ∼0.7- to ∼1.2-fold that of the parental cell line (Fig. 1-Supplement 2 A). Wild-type clones and bortezomib-resistant cell lines treated with taxol had LD50s ranging from ∼0.6- to ∼1.9-fold that of the parental cell line (Fig. 1-Supplement 2 C and D). Finally, treating cells with mitoxantrone, we found that the wild-type clones had LD50s ∼0.6- to ∼3.1-fold that of the parental line (Fig. 1-Supplement 2 E) and most of the bortezomib-resistant clones had similar LD50s (∼0.7- to ∼2.7-fold that of the parental line) (Fig. 1-Supplement 2 F). The exception was BZ06, which had an LD50 nearly 14-fold higher than the wild-type parental line. Since BZ06 did not have reduced sensitivity to taxol (∼0.6-fold that of the parental line) we do not suspect multidrug resistance to be the source of this mitoxantrone resistance. Together, these data indicate that while there is variability in the responses to different treatments, none of the tested cell lines had the expected features of multidrug resistance.

We next applied the Cell Painting assay to these drug-sensitive and -resistant cell lines. Cell Painting captures signal in five imaging channels from six fluorescent dyes that stain cells for eight cellular components including mitochondria, actin, Golgi, plasma membrane, cytoplasmic RNA, nucleoli, endoplasmic reticulum, and DNA (Fig. 1 B) (Bray et al., 2016). With these images, we used CellProfiler (Stirling et al., 2021) to extract single-cell morphological features from individual cells. The signal from each of the five channels was analyzed in the nucleus, cytoplasm, and total cell and characterized based on features (object parameters) such as signal intensity, shape of the object, texture of the staining pattern, etc. yielding a total of ∼3500 features. These cellular features were combined and analyzed on a per well basis and then compared across cell lines and experimental conditions to determine whether morphological features of drug sensitivity could be reliably detected in untreated cells.

of the parental line) we do not suspect multidrug resistance to be the source of this mitoxantrone resistance. Together, these data indicate that while there is variability in the responses to different treatments, none of the tested cell lines had the expected features of multidrug resistance.

We next applied the Cell Painting assay to these drug-sensitive and -resistant cell lines. Cell Painting captures signal in five imaging channels from six fluorescent dyes that stain cells for eight cellular components including mitochondria, actin, Golgi, plasma membrane, cytoplasmic RNA, nucleoli, endoplasmic reticulum, and DNA (Fig. 1 B) (Bray et al., 2016). With these images, we used CellProfiler (Stirling et al., 2021) to extract single-cell morphological features from individual cells. The signal from each of the five channels was analyzed in the nucleus, cytoplasm, and total cell and characterized based on features (object parameters) such as signal intensity, shape of the object, texture of the staining pattern, etc. yielding a total of ∼3500 features. These cellular features were combined and analyzed on a per well basis and then compared across cell lines and experimental conditions to determine whether morphological features of drug sensitivity could be reliably detected in untreated cells.

### Morphology signature of bortezomib resistance distinguishes multiple sensitive versus resistant clones

We first examined whether there were any clear qualitative morphological differences between wild-type and bortezomib-resistant cell lines and chose the wild-type polyclonal parental cell line, wild-type clones WT01-WT05, and bortezomib-resistant clones A, E, and BZ01-BZ05 for these initial studies. We treated cells with 0.1% DMSO (to allow for comparison with future experiments using drug-treated cells) and performed Cell Painting, staining fixed HCT116 cells and imaging as per the published protocol (Bray et al., 2016).

Imaging revealed cellular heterogeneity within each cell line as well as between cell lines with similar bortezomib sensitivities (Fig. 2 A and Fig. 2-Supplement 1). This heterogeneity obscured any potential differences between bortezomib-sensitive and -resistant cell lines and prevented us from qualitatively distinguishing wild-type from bortezomib-resistant clones by eye, confirming the need for high-content quantitative analysis.

**Figure 2.**
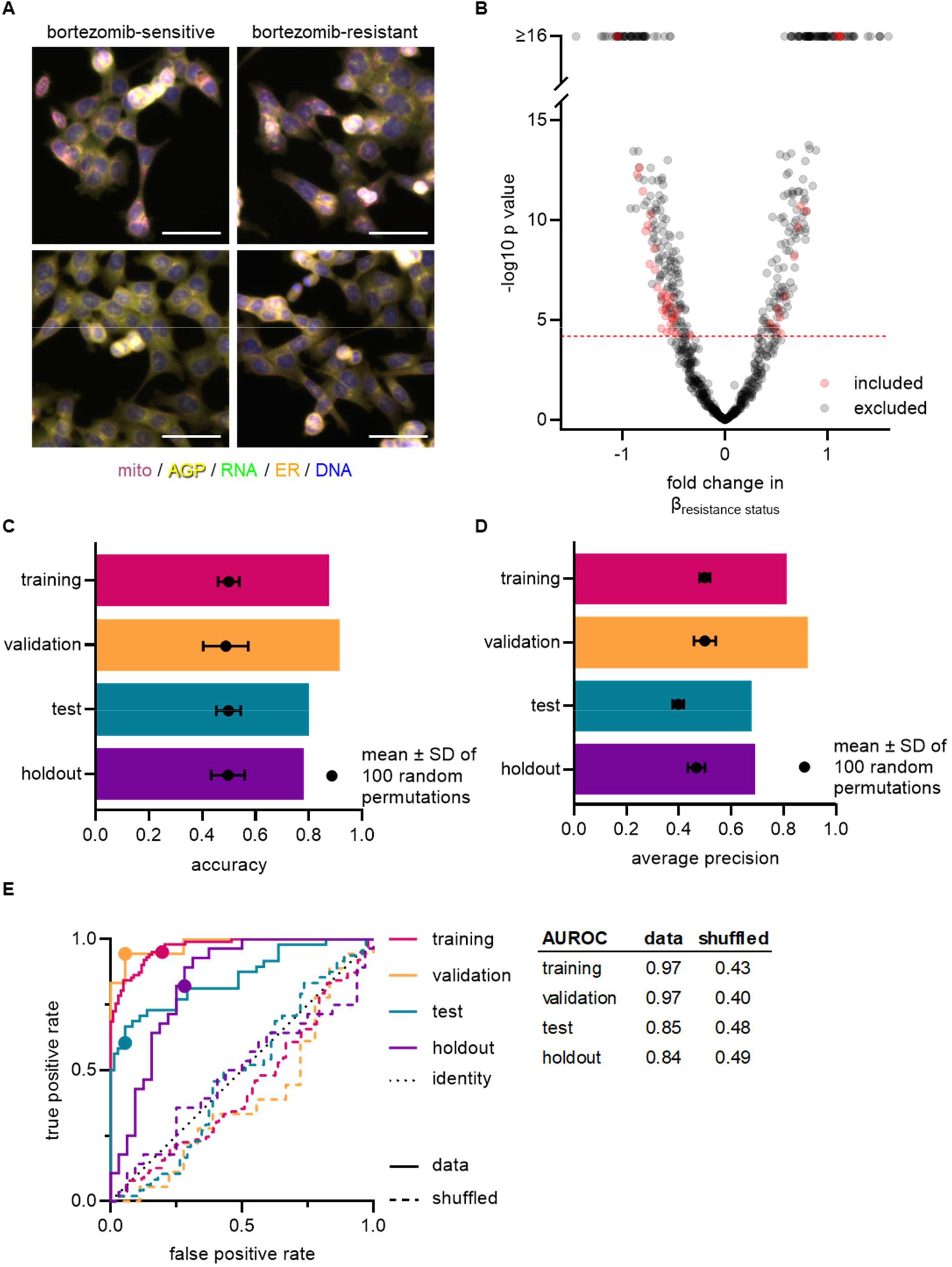
Cell morphology reveals a signature of bortezomib resistance. **(A)** Representative fixed fluorescence microscopy images of two bortezomib-sensitive (WT02 and WT03) and two bortezomib-resistant (BZ02 and BZ03) clones stained and imaged as per the Cell Painting protocol. Channels are labeled as mito (mitochondria; magenta), AGP (actin, golgi, plasma membrane; yellow), RNA (ribonucleic acid; green), ER (endoplasmic reticulum; orange), and DNA (deoxyribonucleic acid; blue). See Figure 2-Supplement 1 for single channel images. Scale bars, 50 μm. **(B)** Volcano plot of the variability of morphological features (β) by resistance status. Y-axis -log_10_p values are from Tukey’s HSD (see Methods). Red circles are features included in the final signature of resistance and gray circles are features excluded from the final signature. Features above the red dashed line (-log_10_[0.05/number of unique features]) were considered significantly varying and those that had not been excluded as technical variables (Fig. 2-Supplement 3) were included in the signature of bortezomib resistance. **(C)-(E)** Evaluations of the Bortezomib Signature on the training (magenta), validation (orange), test (teal), and holdout (purple) datasets. Bar graphs showing the (C) accuracy and (D) average precision of the Bortezomib Signature when characterizing the resistance status of cell lines. Symbols and error bars are the means of the shuffled data ± SD. (E) ROC curves for the performance of the Bortezomib Signature on the indicated dataset (solid line) or its shuffled counterpart (dashed line). Colored points are the corresponding false positive and true positive rates at the absolute minimum thresholds for each respective dataset. Black dotted line is the identity line where false positive rate = true positive rate. AUROC values reported for data and shuffled data. See Fig. 2-Supplement 8 for breakdown of profiles and experiments per dataset.

We then pre-processed profiles to remove low-variance and highly correlated features, and population-averaged single cell measurements at the well level (see Methods). The morphological profiles of wild-type and bortezomib-resistant cells did not cleanly distinguish cell lines based on bortezomib sensitivity (Fig. 2-Supplement 2 A). This was the case even for a short, 4-hour treatment with 7 nM bortezomib (Fig. 2-Supplement 2 B), further indicating the subtlety of any morphological difference between bortezomib-sensitive and -resistant cells and the need for further feature refinement.

Each observed morphological measurement results from a combination of both technical and biological variables. It is therefore important to control and test for technical variables as these can confound subtle signatures that would otherwise dominate the morphological profiles of the cells being analyzed. Using wild-type clones WT01-05 and bortezomib-resistant clones BZ01-05 to quantify and reduce the impact of technical variables, we fit a linear model to each morphological feature adjusting for technical variables (experimental run/batch, incubation time, cell count/density, clone ID) and biological variables (resistance status) (see Methods). We then discarded morphological features with variances that correlated with experimental run (batch), incubation time (4 or 13 hours with 0.1% DMSO), cell density, or those features that varied between two or more pairs of wild-type clones (clone ID) (Fig. 2-Supplement 3 A-E). Of the remaining morphological features, we only considered those that varied based on the resistance status of a cell line (Fig. 2 B). This resulted in 45 morphological features that significantly contributed to cells’ bortezomib drug sensitivity (Fig. 2-Supplement 4). We used these 45 features to compute a resistance score or “Bortezomib Signature” for each cell line based on the direction-sensitive ranking method for phenotype analysis, singscore (Foroutan et al., 2018). With the exception of some texture-based features, the Bortezomib Signature features were largely independent, displaying low pairwise correlation, which may implicate a more nuanced phenotype of drug resistance and explain why detecting resistance by eye was so challenging (Fig. 2-Supplement 5).

Anticipating well location as a possible technical artifact, we plated our cells in a repeating serpentine pattern, ensuring that each cell line would be imaged in multiple locations across each plate (Fig. 2-Supplement 6 A). We found that the pattern of Bortezomib Signatures corresponded to the cell identity plate layout (Fig. 2-Supplement 6 B), indicating that the well position for each cell line was not strongly contributing to its Bortezomib Signature. In addition, we found that the Bortezomib Signature correlated with resistance status of cell lines and not technical variables (Fig. 2-Supplement 7). These data suggest that our analysis pipeline and signature building process minimized technical artifacts.

To evaluate the performance of our Bortezomib Signature, we used machine learning best practices, separating our data into training, validation, test, and holdout datasets (Fig. 2-Supplement 8; see Methods). The data used to create the Bortezomib Signature, which included well-based morphological profiles from clones WT01-05 and BZ01-05, was designated as the training dataset, which we used to build the Bortezomib Signature initially. The validation dataset was composed of profiles from clones WT01-05 and BZ01-05 that were not used to generate the Bortezomib Signature but were collected on the same plates as the profiles used for the training dataset. The test dataset was composed of profiles from the wild-type parental cell line and bortezomib-resistant clones A and E (none of these lines were included in training), and these profiles were also collected on the same plates as those used for the training dataset. The holdout dataset was a separate plate and contained wild-type parental cells, clones WT01-05, and bortezomib-resistant clones A, E, and BZ01-05. These datasets allowed us to test generalizability across clones and plates for the trained Bortezomib Signature.

We found the Bortezomib Signature could predict whether a cell line was bortezomib-resistant or bortezomib-sensitive (Fig. 2 C and D and Fig. 2-Supplement 9 A-D). We called the prediction bortezomib-resistant if the Bortezomib Signature was greater than zero and bortezomib-sensitive for a Bortezomib Signature less than zero. In the training dataset, the Bortezomib Signature correctly characterized cell lines as either sensitive or resistant to bortezomib 88% of the time with an average precision of 81%. The signature performed similarly well in the validation dataset (wells not included in the training dataset), with an accuracy of 92% and an average precision of 89%, as would be expected given that the validation dataset included the same clones and same plates used for the training dataset. In the test dataset, composed solely of wild-type parental cells and bortezomib-resistant clones A and E, the Bortezomib Signature had an accuracy of 80% and an average precision of 68%. Similarly, in the holdout dataset the Bortezomib Signature had an accuracy of 78% and an average precision of 69%. Although the Bortezomib Signature did not perform as well in the test and holdout datasets as it did in the training and validation datasets, this was expected given that the test dataset included the polyclonal wild-type parental cell line and two previously isolated bortezomib-resistant clones while the holdout dataset was collected on a single, unique plate. However, the Bortezomib Signature performed better than random chance in all testing conditions, as demonstrated by comparison with the mean values for the randomly shuffled data, and as reflected in receiver operating characteristic (ROC) curves, which describe the classification trade-off between true positive and false positive rates in predicting bortezomib-resistance (Fig. 2 E).

### Bortezomib Signature is specific to bortezomib over other ubiquitin-proteasome system inhibitors

We next tested whether the Bortezomib Signature is specific to the drug bortezomib or more broadly to the UPS. To test this, we performed Cell Painting on HCT116 cell lines that were resistant to either ixazomib (another proteasome inhibitor that targets the PSMB5 subunit) or CB-5083 (a p97 inhibitor that acts upstream of the proteasome). If our Bortezomib Signature was a general signature of UPS-targeting drug resistance, we would expect our signature to perform equally well at characterizing the drug sensitivity of bortezomib-, ixazomib-, and CB-5083-resistant cell lines. Our Bortezomib Signature performed better than chance at identifying ixazomib-resistant and CB-5083-resistant cell lines (Fig. 3 A), correctly identifying four of five ixazomib-resistant clones (Fig. 3 B) and three of five CB-5083-resistant clones (Fig. 3 C). However, the areas under the ROC curves for these clones (0.63 and 0.60, respectively) were lower than those observed for any of our bortezomib-resistant datasets and many of these Bortezomib Signatures, particularly those for CB-5083-resistant clones, landed within the range of randomly permuted data. These results suggest that our Bortezomib Signature is not a general signature of UPS-targeting and is relatively specific to bortezomib or the proteasome.

**Figure 3.**
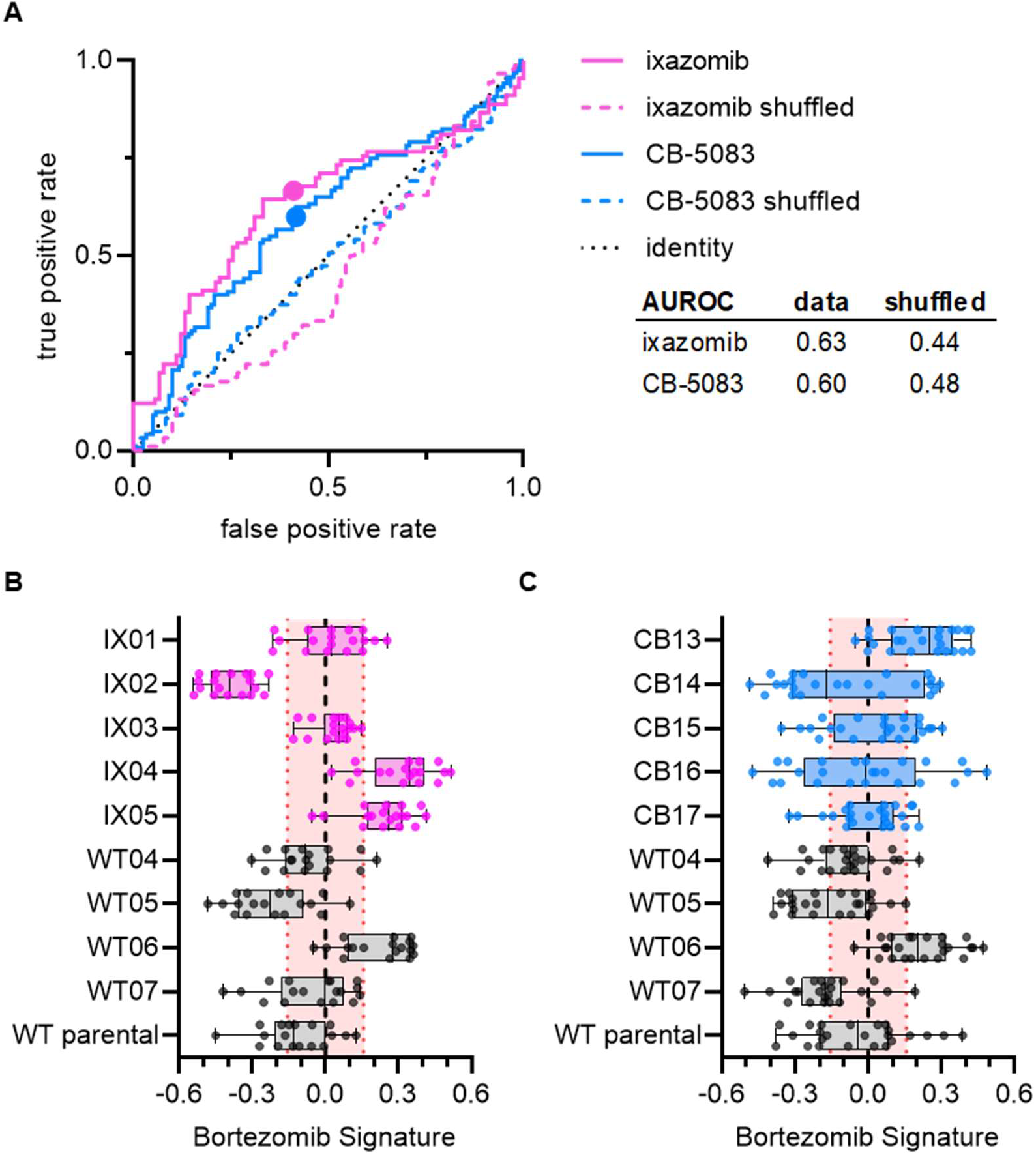
Signature of bortezomib resistance is specific to drug and pathway. **(A)** ROC curves for ixazomib-resistant (magenta) and CB-5083-resistant (blue) experimental data. Colored solid lines are the actual data while colored dashed lines are the shuffled data for each set of clones. Colored points are the corresponding false positive and true positive rates at the absolute minimum thresholds for each respective cell type. Black dotted line is the identity line where false positive rate = true positive rate. AUROC reported for the data and shuffled data. Box plots of Bortezomib Signatures for **(B)** ixazomib-resistant and wild-type cell lines (*n* = 18 profiles, 3 independent experiments) and **(C)** CB-5083-resistant and wild-type cell lines (*n* = 24 profiles, 4 independent experiments). Plots show individual points, range (error bars), 25th and 75th percentiles (box boundaries), and median. Dashed vertical black line is Bortezomib Signature = 0, dashed vertical red lines are the 95% confidence interval for Bortezomib Signatures of 1000 random permutations of the data.

### Bortezomib Signature characterizes bortezomib sensitivity of cell lines not included in the training dataset

To test whether our Bortezomib Signature could correctly characterize the bortezomib sensitivity of cell lines not included in the training dataset, we imaged an entirely new set of wild-type (WT10, WT12-WT15) and bortezomib-resistant clones (BZ06-BZ10) using the Cell Painting protocol. Overall, the Bortezomib Signature performed well, with an AUROC of 0.75, compared to 0.55 for the shuffled data (Fig. 4 A). Our Bortezomib Signature correctly characterized the bortezomib sensitivity of our wild-type bortezomib-sensitive parental line and our bortezomib-resistant clones A and E, which we included as controls (Fig. 4 B), as well as four of five bortezomib-resistant clones and three of five wild-type clones not included in the training dataset (Fig. 4 C). In addition, the majority of these Bortezomib Signatures landed outside the range of randomly permuted data. These results indicate the drug-specificity of our signature and suggest that this Bortezomib Signature has the potential to identify bortezomib-resistant cell lines based on the intrinsic morphological features of untreated cells.

**Figure 4.**
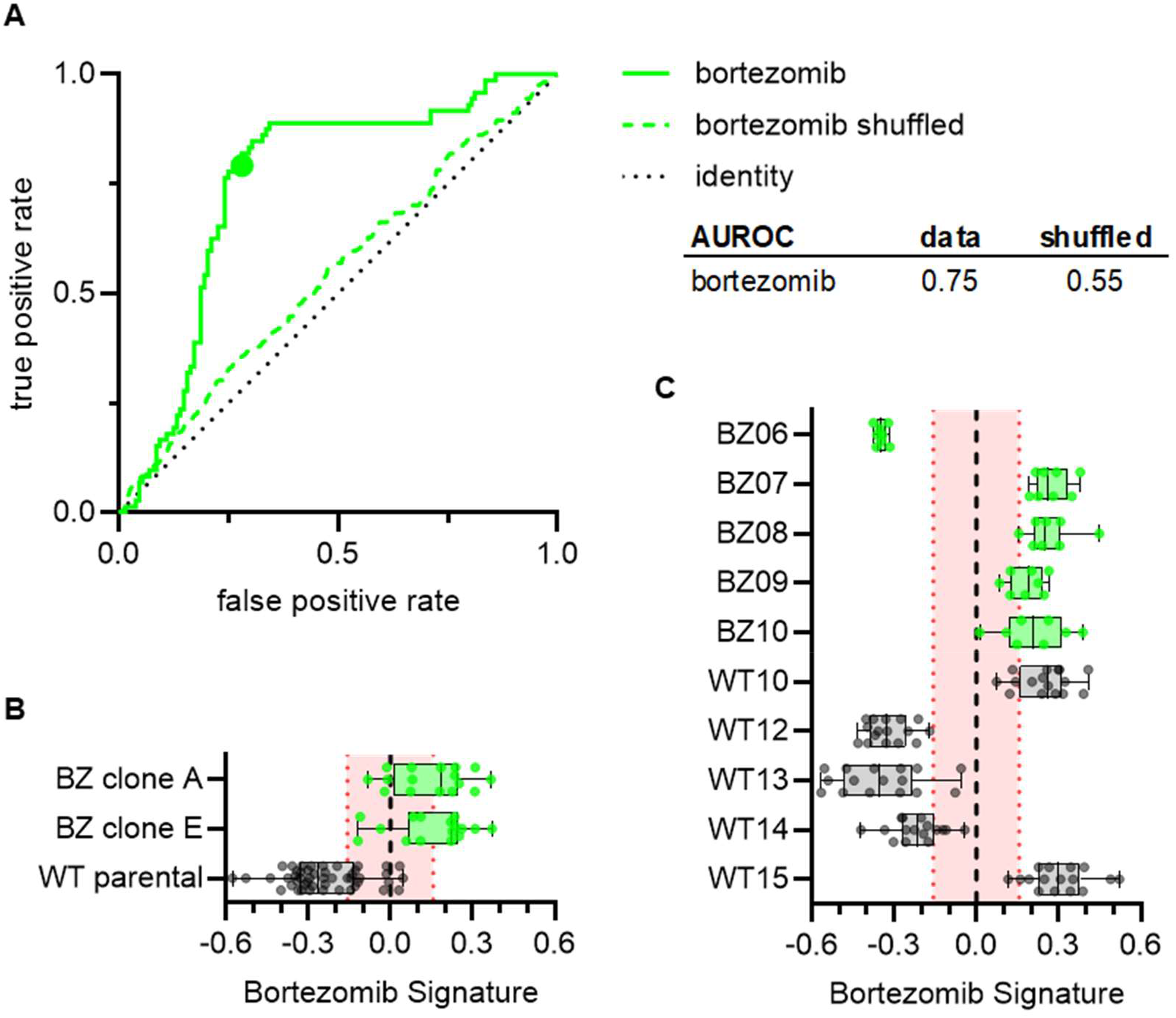
Bortezomib Signature correctly predicts bortezomib sensitivity of cell lines not included in the training dataset. **(A)** ROC curve for the performance of the Bortezomib Signature on the cell lines in (B) and (C) (solid line) and shuffled data (dashed line). Colored point is the corresponding false positive and true positive rate at the absolute minimum threshold. Black dashed line is the identity line where false positive rate = true positive rate. AUROC reported for the data and shuffled data. **(B)** Box plots of Bortezomib Signatures for bortezomib-resistant clones A and E (*n* = 16 profiles each) and wild-type parental cells (*n* = 48 profiles). **(C)** Box plots of Bortezomib Signatures for wild-type clones WT10, WT12-15 (*n* = 16 profiles each) and bortezomib-resistant clones BZ06-BZ10 (*n* = 8 profiles each). Plots show individual points, range (error bars), 25th and 75th percentiles (box boundaries), and median. Dashed vertical black line is Bortezomib Signature = 0, dashed vertical red lines are the 95% confidence interval for Bortezomib Signatures of 1000 random permutations of the data.

## DISCUSSION

We used Cell Painting, a high-throughput and high-content image acquisition and analysis assay, as a target-independent method to capture the morphological profiles of untreated cells that were either sensitive or resistant to the ubiquitin-proteasome system (UPS)-targeting anticancer drug, bortezomib. After processing profiles to reduce the impact of technical variables, we generated a signature of bortezomib resistance and characterized the performance of this signature using machine learning best practices. This Bortezomib Signature correctly predicted the bortezomib treatment response of seven out of ten cell lines not included in the training dataset and was specific to the drug under investigation, in this case bortezomib, even as compared to cells that were resistant to other drugs targeting the UPS. Our work demonstrates that there are intrinsic morphological features of drug resistance in cells that can be identified using Cell Painting and provides a reproducible pipeline for generating morphological signatures of drug resistance.

The Bortezomib Signature’s performance was not perfect; it mischaracterized three clones not included in the training dataset. Interestingly, one of the misidentified clones (BZ06) had reduced sensitivity to mitoxantrone as well as bortezomib. Given the considerable genetic heterogeneity in this mismatch repair-deficient HCT116 cell line (Glaab and Tindall, 1997; Umar et al., 1994), it is possible that these mischaracterized cell lines have orthogonal mechanisms of resistance or unrelated mutations contributing to their morphological profiles. Targeted sequencing of the PSMB5 proteasome subunit in bortezomib-resistant clones may yield clues to the origins of these misidentifications, as multiple mutations have been identified in bortezomib-resistant cell lines (Wacker et al., 2012). Determining the underlying reason for the misidentification of wild-type cell lines would require more comprehensive sequencing.

Our Bortezomib Signature performed better at identifying bortezomib-resistant cell lines compared to ixazomib-resistant cell lines, and better at identifying ixazomib-resistant cell lines compared to CB-5083-resistant cell lines. All three drugs broadly target the UPS, however bortezomib and ixazomib both target the same subunit of the proteasome, albeit with different potentially non-overlapping spectrums of off-targets (Kupperman et al., 2010). Our data therefore suggest that the Bortezomib Signature is specific to the drug bortezomib, and not proteasome inhibition broadly or simply a general signature of UPS-targeting drug resistance.

This work has shown potential for morphological profiling with Cell Painting to characterize drug sensitivity in untreated cells, having generated a robust signature of resistance to bortezomib, a drug with a high failure rate in treating cancer. Our results indicate that different mechanisms of bortezomib resistance may be generating distinct morphological profiles; with larger and broader training data, it may be possible to identify signatures for multiple mechanisms of bortezomib resistance as well as signatures of resistance to other drugs. An important step will be determining whether this method can be extended to patient samples where identifying intrinsic drug resistance in cells prior to treatment has the potential to improve targeted cancer therapy. We expect that further refinement might develop Cell Painting as a tool for identifying drug-resistant cells, perhaps even guiding therapeutic strategies to overcome intrinsic resistance.

## MATERIALS AND METHODS

### Cell culture

HCT116 cells (CCL-247; ATCC), also referred to as HCT116 wild-type parental cells, were maintained in McCoy’s 5A Medium (Gibco) supplemented with 10% (v/v) FBS (Sigma) and cultured at 5% CO2 and 37°C. Bortezomib-resistant, ixazomib-resistant, and CB-5083-resistant clonal cell lines were isolated as previously described (Wacker et al., 2012). Briefly, cells were plated in 150mm dishes and grown in the presence of approximately the LD90 of the desired drug until the majority of cells died. The locations of single surviving cells were marked and the colonies that expanded were isolated using cloning rings. HCT116 wild-type clonal cell lines were isolated by dilution into 96-well plates and wells containing single cells that expanded into colonies were selected. Bortezomib-resistant clones A and E were provided by the Kapoor laboratory having been previously published (Wacker et al., 2012).

### Proliferation assays

Cell proliferation was evaluated using an Alamar Blue assay (O’Brien et al., 2000). Briefly, cell lines were plated in duplicate or triplicate in sterile 96-well Clear Microplates (Falcon) under described culture conditions, with 1000 cells in 100 μL per well and allowed to adhere overnight. After cells attached to the plate, 50 μL of media containing drug was added to each well. The final DMSO concentration was 0.1% for all wells, including three wells with media only as background measurements. Plates were incubated for 72 hours at 5% CO2 and 37°C before adding Alamar Blue (resazurin sodium salt, final concentration 50 μM). Cells were incubated with Alamar Blue for 3-4 hours and then imaged with a Synergy Neo plate reader using excitation: 550 nm and emission: 590 nm (Agilent). The average plate background (media only with 0.1% DMSO) was subtracted from the average fluorescence for each condition and the resulting value was normalized by dividing by the background-subtracted value for each condition’s control (cells treated with 0.1% DMSO). With the data from our proliferation assays, we calculated the median lethal dose (LD50) for each of our drugs of interest by fitting data of normalized growth vs. log[drug concentration] to a sigmoidal dose-response curve using GraphPad Prism (v.9.2.0) (Fig. 1-Supplement 1 D) and then determined the dose at which 90% of cells would be expected to die (LD90).

### Cell Painting

High-throughput imaging was performed according to the published Cell Painting protocol (Bray et al., 2016). HCT116 cells were plated at concentrations of 2.5 or 5 × 10^3^ cells/mL in 96-well glass-bottomed tissue culture dishes (Greiner Bio-One) and allowed to adhere for 48-72 hours prior to fixation. At either 4 or 13 hours prior to fixation, cells were treated with either 0.1% DMSO or 7 nM bortezomib and 30 min prior to fixation cells were treated with MitoTracker Deep Red (500 nM, Invitrogen). 16 % paraformaldehyde (EMS) was added to each well for a final concentration of 3.2% and cells were fixed in the dark at room temperature for 20 minutes. Wells were washed with HBSS (Invitrogen), permeabilized with 0.1% Triton-X for 15 minutes, and then washed twice with HBSS before incubating with staining solution (5 U/mL phalloidin AF568 [Invitrogen], 100 μg/mL concanavalin A AF488 [Invitrogen], 5 μg/mL Hoechst 33342 [ThermoFisher or Invitrogen], 1.5 μg/mL wheat-germ agglutinin AF555 [Invitrogen], 3 μM SYTO14 Green [Invitrogen], and 1% bovine serum albumin [BioWorld] in HBSS) in the dark for 30 minutes. Wells were then washed twice with HBSS and imaged using an ImageXpress high-content imaging system (Molecular Devices) with a 20x 0.45 NA S Plan Fluor ELWD objective (Nikon) and captured with a Zyla 5.5 sCMOS detector (Andor Technology). Each well was imaged at 12-17 non-overlapping sites in five channels using Semrock filters (mito: Cy5-4040B-NTE-ZERO, AGP: TxRed-4040C-NTE-ZERO, RNA: Cy3-4040C-NTE-ZERO, ER: FITC-3540C-NTE-ZERO, and DNA: DAPI-5060C-NTE-ZERO).

### Image data processing

We used CellProfiler versions 3.1.8 and 3.1.9 (McQuin et al., 2018) to perform the standard processing pipeline of illumination correction, single cell segmentation, and morphology feature extraction. We performed per-plate illumination correction to adjust for uneven background intensity that commonly impacts microscopy images. We also developed per-plate analysis pipelines for single cell segmentation and feature extraction. We extracted 3,528 total cell morphology features from all 25,331,572 cells we generated in this experiment. The 3,528 features represent stain intensities, stain co-localization, textures, areas, and other patterns extracted from all five imaging channels and different segmentation objects (nuclei, cytoplasm, total cells). Feature details are described in the documentation for CellProfiler (https://cellprofiler-manual.s3.amazonaws.com/CellProfiler-3.1.9/help/output_measurements.html).

Following feature extraction, we applied an image-based analysis pipeline to generate our final analytical set of treatment profiles (Caicedo et al., 2017). We first used cytominer-database to ingest all single-cell, per-compartment CellProfiler output files (comma separated) to clean column names, confirm integrity of CellProfiler output CSVs, and output single-cell SQLite files for downstream processing. Next, we used pycytominer (github hash c1aa34b641b4e07eb5cbd424166f31355abdbd4d) for all image-based profiling pipeline steps. In the first step, we median aggregated all single cells to form well-level profiles (Way et al., 2022). Next, we performed a step called annotation, which merges the consistent platemap metadata with the well-level profiles. Third, we performed standard z-score normalization to ensure all features are measured on the same scale with zero mean and unit variance. Lastly, we performed feature selection, which removed features with low variance, high correlation (>0.9 Pearson correlation), features with missing values, features on our blocklist (Way, 2020), and features with outliers greater than 15 standard deviations, which we suspected were measured in error. For developing our final analytical datasets (see next section) we performed normalization within each plate but performed a combined feature selection across all plates per analytical dataset using the same procedures described previously, which resulted in 782 features. We applied the same pipeline uniformly across all plates. We did not detect large differences in variance that could be attributed to well position and batch and therefore did not apply batch effect correction. Our full image data processing pipeline is publicly available at https://github.com/broadinstitute/profiling-resistance-mechanisms (Way et al., 2023).

### Constructing the resistance signature

After processing all images and forming normalized and feature selected profiles per well, we performed several additional analyses to explore the results and discover a morphology profile of bortezomib resistance. We performed initial comparisons of morphological profiles using Morpheus (https://software.broadinstitute.org/morpheus) to create similarity matrix heatmaps.

We aimed to discover a generalizable signature of bortezomib resistance from the normalized profiles. Our approach was to identify features that were significantly different by resistance status and not significantly impacted by technical covariates. To do so, we carefully constructed datasets for training and evaluating signature performance (Fig. 2-Supplement 8). To generate our training dataset, we selected a set of six plates consisting of five wild-type and five bortezomib-resistant clones that we collected on three different days, which showed high within-replicate reproducibility (data not shown). A seventh plate was held-out from signature generation in order to analyze generalizability between plates (holdout dataset). We evaluated the signature in five scenarios: 1) clones held-out on the same plates used to generate the training dataset (validation dataset, Fig. 2-Supplement 9 B), 2) wild-type parental cells and clones with confirmed PSMB5 mutations known to confer resistance to bortezomib (test dataset, Fig. 2-Supplement 9 C) (Wacker et al., 2012), 3) clones held-out on a separate plate (holdout dataset, Fig. 2-Supplement 9 D), 4) clones selected to be resistant to other drugs (ixazomib and CB-5083, Fig. 4), and 5) bortezomib-resistant clones not included in the training dataset (Fig. 3). All cells on these plates were incubated with 0.1% DMSO for either 4 or 13 hours.

Using data from the ten clones in our training dataset (20-21 replicates per clone, see Fig. 2-Supplement 8), we fit two linear models for all 782 CellProfiler features (post normalization and feature selection) to discover features that varied strongly with technical variants (batch, cell count, incubation time, or clone ID) and features that varied strongly with resistance status (wild-type or resistant). In the first linear model, we quantified the per feature variance contribution of resistance status (β_resistance status_), batch (β_batch_), incubation time (β_incubation time_), and clone (β_clone ID_) to each CellProfiler feature (Y_j_) where *ε* is the error term:

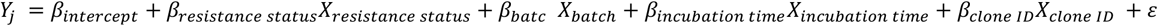

Fitting this model produced a goodness of fit R^2^ value per feature and individual beta coefficients per covariate. Furthermore, we calculated a Tukey’s Honestly Significant Difference (Tukey’s HSD) post hoc test per model to determine which specific categorical covariate comparison contributed to a significant finding and to control for within-covariate-group multiple comparisons through a family-wise error rate (FWER) adjustment that accounts for different within-group sizes (e.g. three different batches in the comparison, two different resistance statuses, etc.)(Tukey, 1949).

Separately, we fit another linear model on continuous features to adjust for features that were significantly impacted by well confluence (β_cell count_) as it is expected that dense wells will impact certain morphology features, which we want to avoid in our resistance signature:

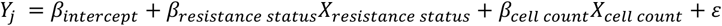

By fitting these models, we quantified the variance contribution of four technical covariates (incubation time, batch, clone ID, and cell count) and our biological variable of interest (resistance status), and, based on the first linear model, we have knowledge of which specific group comparisons were significant in each category (via Tukey’s HSD). We further refined the signature by filtering features that did not pass a Bonferonni adjusted alpha threshold calculated across all 782 features (0.05 / 782 = 6.4×10^−6^).

We next applied a specific exclusion criterion to specifically isolate features that contributed to resistance status. We excluded features that were significantly different across incubation times, batches, and cell counts. We also excluded features that were different within clone type (features varying between two or more wild-type clones) to reduce the contribution of features that may mark generic inter-cell line differences nonspecific to resistance status. This procedure resulted in a total of 45 features that were significantly different by resistance status and not significantly impacted by any of the technical covariates we considered. Of the 45 features, 14 had higher values in resistant clones and 31 had lower values in resistant clones (Fig. 2-Supplement 4).

### Applying the signature

We used the *singscore* method (Foroutan et al., 2018) to characterize individual profiles of different cell lines as either bortezomib-sensitive or bortezomib-resistant. *Singscore* is a rank-based method that was originally developed to analyze the direction and significance of previously defined molecular signatures on transcriptomic data. The method calculates a two-part signature for each direction list (14 up and 31 down) and calculates an internal rank per profile of how highly ranked and lowly ranked each of the up and down features are, respectively. The method then adds the up and down rank scores to form a total *singscore* per sample, which ranges between -1 and 1 and represents a rank-based normalized concordance score that can be directly compared across profiles that may have been normalized differently. Therefore, the score is robust to outliers and different normalization procedures. In addition to calculating the *singscore* per sample, we also calculated *singscore* with 1,000 random permutations, in which we randomly shuffled feature rankings to derive a range in which a sample may be scored by chance.

### Signature evaluation

We used several metrics to evaluate signature quality across five different evaluation scenarios (validation, test, holdout, other UPS-targeting drugs, and clones not included in the training dataset). Because we are measuring a binary decision in a balanced dataset (roughly the same amount of positive as negative classes), we used accuracy (total correct / total chances) to quantify performance. We also calculated mean average precision using sci-kit learn, averaging over samples along the precision recall curve (Pedregosa et al., 2011), which is a measure of separation between the two resistance classes (higher being more separation). We also calculated receiver operating characteristic (ROC) curves and area under the ROC curve (AUROC) using sci-kit learn. AUROC compares the ability to distinguish positive samples across signatures.

## AUTHOR CONTRIBUTIONS

Conceptualization: MEK, AYB, SS, AEC, TMK, GPW

Data curation: MEK, AYB, DRS, BAC, GPW

Formal analysis: MEK, AYB, DRS, YH, GPW

Funding acquisition: AEC, TMK

Investigation: MEK, AYB, DRS, BAC, YH, SS, AEC, TMK, GPW

Methodology: MEK, AYB, DRS, GPW

Project administration: BAC, SS, AEC, TMK, GPW

Resources: SS, AEC, TMK

Software: DRS, BAC, YH, GPW

Supervision: BAC, SS, AEC, TMK, GPW

Validation: MEK, AYB, DRS, GPW

Writing - original draft: MEK, GPW

Writing - review and editing: MEK, AYB, DSR, BAC, YH, SS, AEC, TMK, GPW

## ACKNOWLEDGEMENTS

The authors gratefully acknowledge funding from the Starr Cancer Consortium (112-0039 to TK and AEC), the National Institutes of Health (NIH MIRA R35 GM122547 to AEC, NIH MIRA R35 GM130234 to TMK, NIH NRSA T32 GM066699 to MEK, NIH T32 GM115327 Chemistry-Biology Interface Training Grant to the Tri-Institutional PhD Program in Chemical Biology to AYB), and the National Science Foundation (NSF GRFP 2019272977 to AYB).

