## Supplementary Figures for "High-content microscopy reveals a morphological signature of bortezomib resistance"

*co-senior authors

Figure 1-Supplement 1. HCT116 parental cells were grown in select drugs to isolate drug-resistant clones. 2

Figure 1-Supplement 2. Drug-resistant clones did not display features of multidrug resistance. 4

Figure 2-Supplement 1. Individual channels of bortezomib-sensitive and bortezomib-resistant clones imaged by Cell Painting. 6

Figure 2-Supplement 2. Similarity clustering was insufficient to distinguish wild-type from bortezomib-resistant clones. 8

Figure 2-Supplement 3. Technical variables were controlled for and excluded from analyses. 10

Figure 2-Supplement 4. Select features contributed to the signature of bortezomib resistance. 12

Figure 2-Supplement 5. Bortezomib Signature-contributing features did not universally correlate. 13

Figure 2-Supplement 6. Plating pattern does not strongly correlate with Bortezomib Signature. 14

Figure 2-Supplement 7. Bortezomib Signature identifies resistant clones and not technical variables. 15

Figure 2-Supplement 8. Datasets for Bortezomib Signature generation and initial evaluation. 17

Figure 2-Supplement 9. Bortezomib Signatures from individual cell lines in training, validation, test, and holdout datasets. 18

**Figure 1-Supplement 1. HCT116 parental cells were grown in select drugs to isolate drug-resistant clones**

A

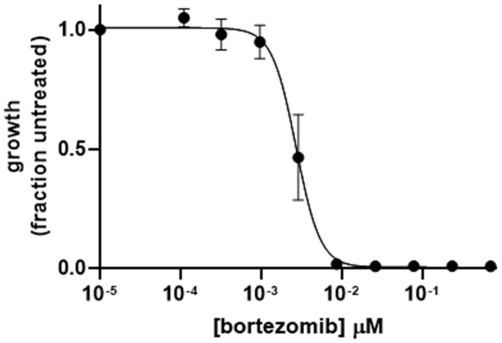


B

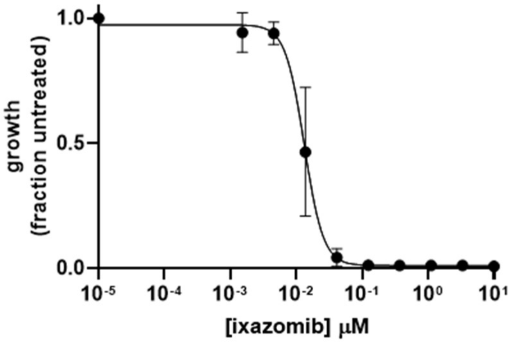


C

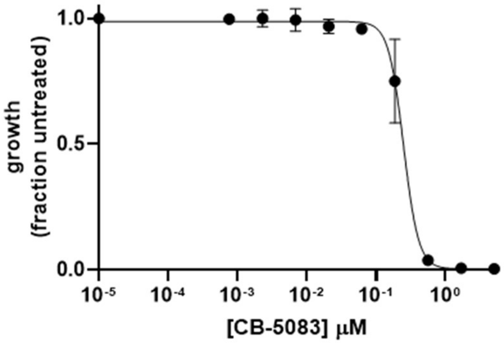


D

| Drug | LD50 |
|---|---|
| bortezomib | ~2.7 nM |
| ixazomib | ~13 nM |
| CB-5083 | ~246 nM |

E

| Designation | Drug-Resistance | LD50 | MDR |
|---|---|---|---|
| HCT116 parental | none | NA | no |
| WT01-05, 10, 12-15 | none | NA | no |
| WT06-07 | none | NA | n.d. |
| BZ01-10 | bortezomib | ~11-35 nM | no |
| *clones A and E | bortezomib | ~10 and ~23 nM | no |
| IX01-05 | ixazomib | n.d. | n.d. |
| CB13-17 | CB-5083 | n.d. | n.d. |

**Figure 1-Supplement 1. HCT116 parental cells were grown in select drugs to isolate drug-resistant clones. (A)-(C)** Proliferation assays of HCT116 parental cells treated with (A) bortezomib, (B) ixazomib, and (C) CB-5083. Growth is measured relative to untreated cells. Mean ± SD. *n* = 3 independent experiments with 3 technical replicates per condition. **(D)** Calculated LD50s from the results of proliferation assays in (A)-(C) using HCT116 parental cells and either bortezomib, ixazomib, or CB-5083. **(E)** Descriptions of the parental and clonal HCT116 cells used in experiments. NA, not applicable; n.d., not determined; * isolated previously (Wacker et al., 2012).

**Figure 1-Supplement 2. Drug-resistant clones did not display features of multidrug resistance**

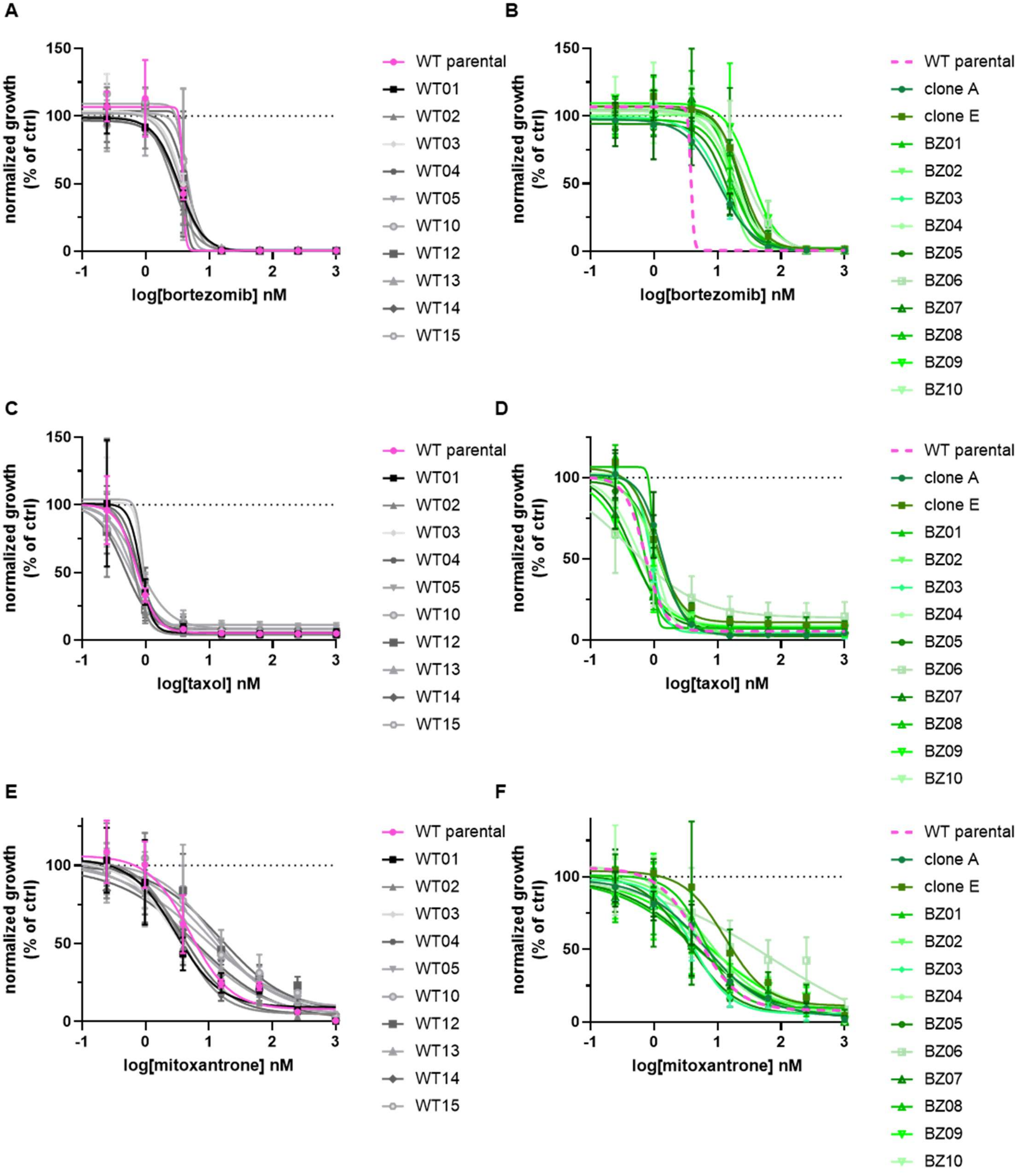

**Figure 1-Supplement 2. Drug-resistant clones did not display features of multidrug resistance.** Proliferation assays of wild-type (left) and bortezomib-resistant (right) HCT116 cell lines. Cells were treated with **(A)-(B)** bortezomib, **(C)-(D)** taxol, or **(E)-(F)** mitoxantrone**.** Dashed magenta line in each bortezomib-resistant graph represents data from the HCT116 parental cell line in the corresponding wild-type graph. Growth is measured relative to untreated control cells (black dotted line). Mean ± SD. *n* = 3 independent experiments per condition.

**Figure 2-Supplement 1. Individual channels of bortezomib-sensitive and bortezomib-resistant clones imaged by Cell Painting**

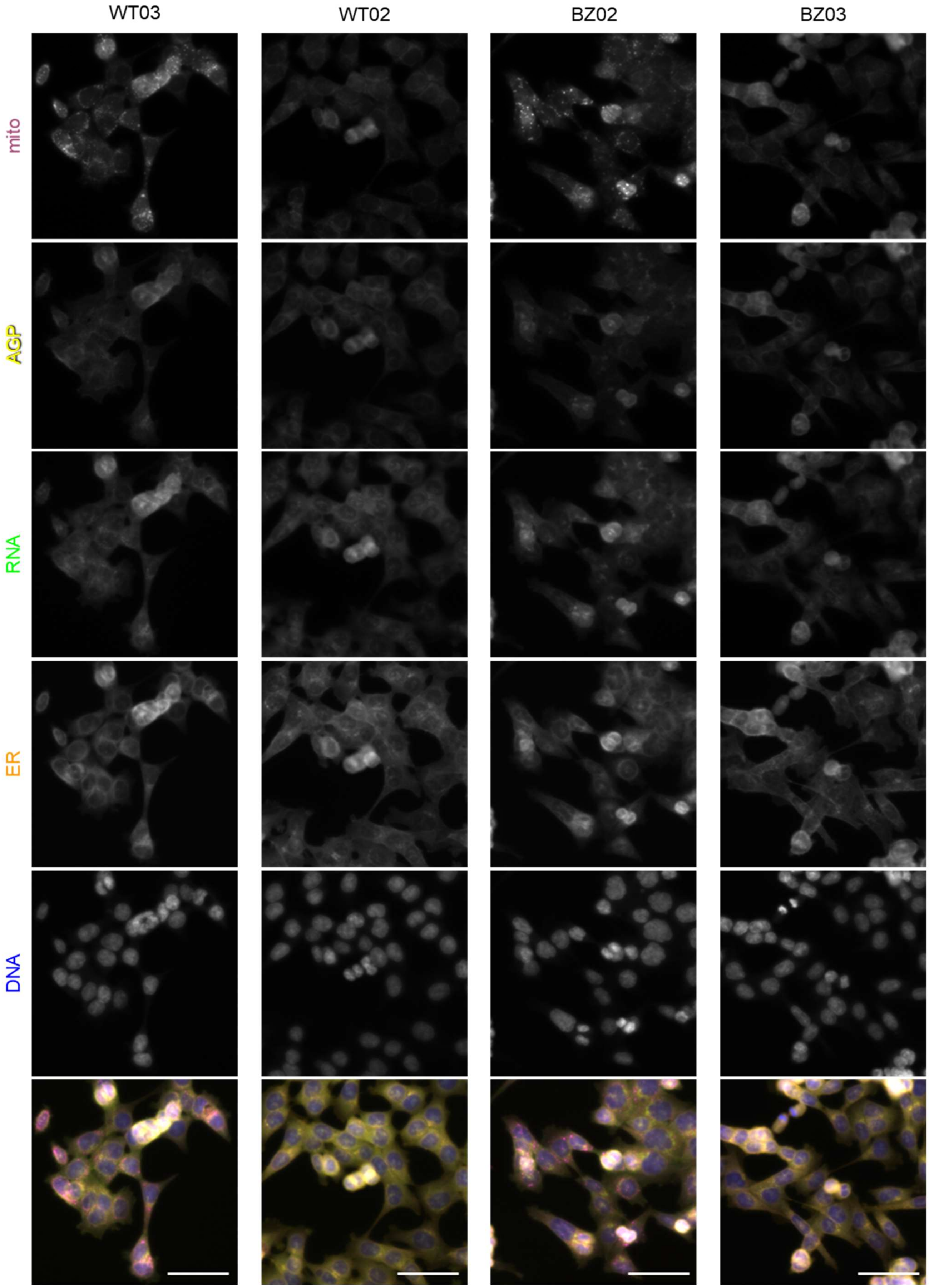

**Figure 2-Supplement 1. Individual channels of bortezomib-sensitive and bortezomib-resistant clones imaged by Cell Painting.** Representative single-channel fixed fluorescence microscopy images of two bortezomib-sensitive (left) and two bortezomib-resistant (right) clones from Fig. 2 A stained and imaged as per the Cell Painting protocol. Channels are labeled as mito (mitochondria; magenta), AGP (actin, golgi, plasma membrane; yellow), RNA (ribonucleic acid; green), ER (endoplasmic reticulum; orange), and DNA (deoxyribonucleic acid; blue). Scale bars, 50 μm.

**Figure 2-Supplement 2. Similarity clustering was insufficient to distinguish wild-type from bortezomib-resistant clones**

**A**

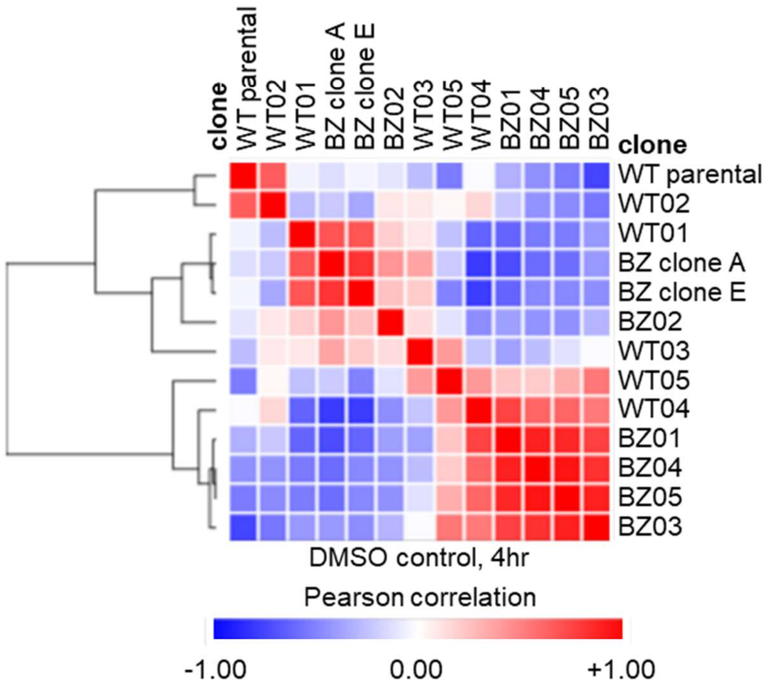


**B**

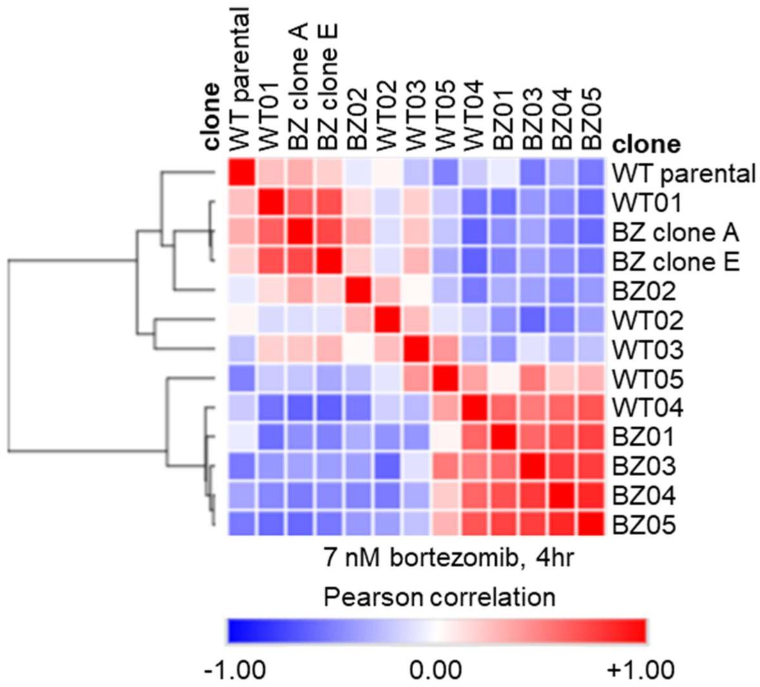

**Figure 2-Supplement 2. Similarity clustering was insufficient to distinguish wild-type from bortezomib-resistant clones.** Similarity matrices of pair-wise Pearson correlation coefficients for morphology profiles of bortezomib-sensitive wild-type cell lines and bortezomib-resistant cell lines treated for 4 hours with either **(A)** 0.1% DMSO or **(B)** 7 nM bortezomib. Dendrograms display hierarchical clustering of pairwise similarity.

**Figure 2-Supplement 3. Technical variables were controlled for and excluded from analyses**

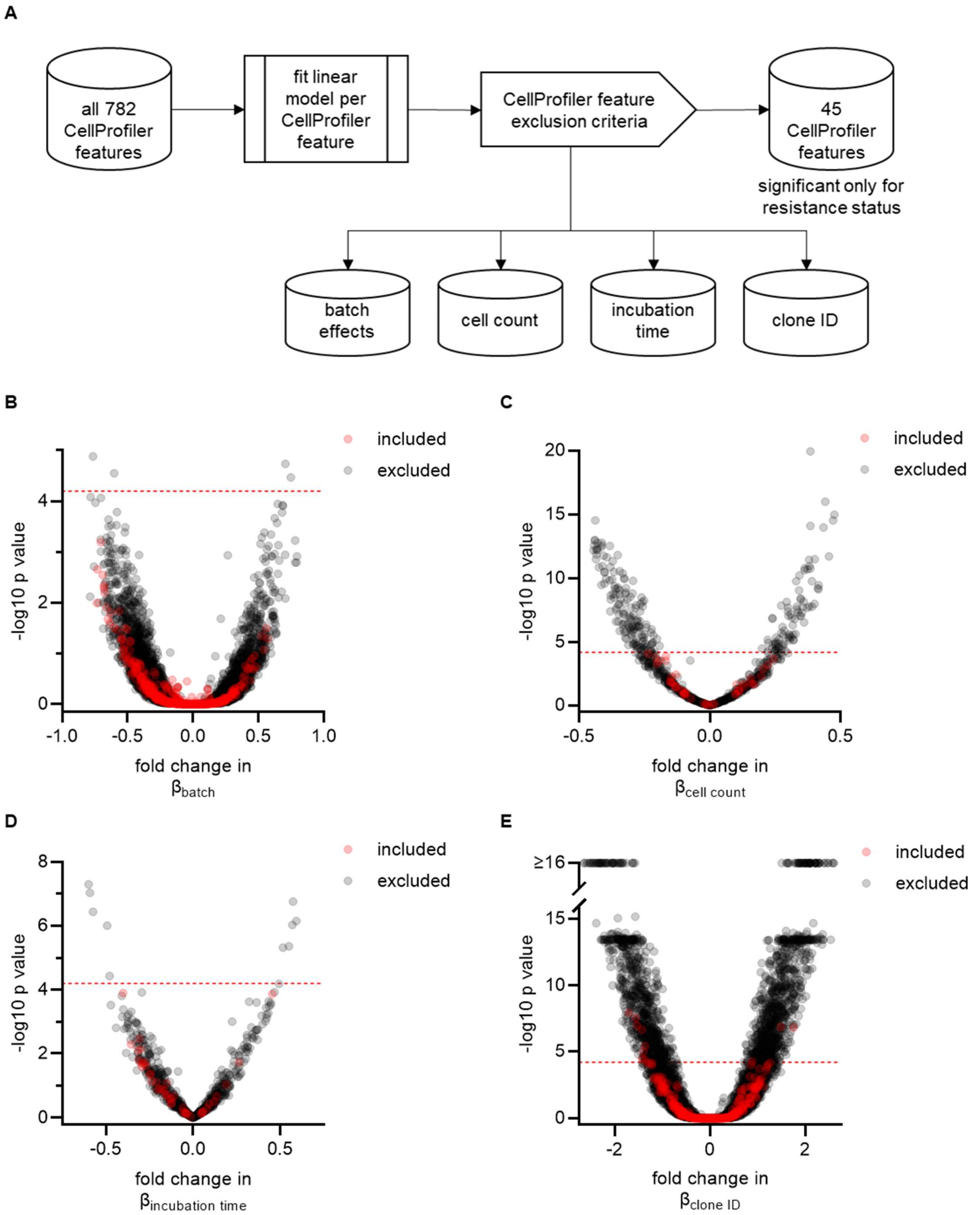

**Figure 2-Supplement 3. Technical variables were controlled for and excluded from analyses. (A)** Model of the workflow to exclude CellProfiler features not related to resistance status. **(B-E)** Volcano plots of the variability of morphological features (β) by (B) batch, (C) cell count, (D) incubation time, and (E) clone ID. Y-axis $-\log_{10}p$ values are from Tukey's HSD (B and D-E) and linear regression analysis (C). Red circles are features included in the final signature of resistance and gray circles are features excluded from the final signature. For technical variables (B-E), features above the red dashed line ($-\log_{10}$[0.05/number of unique features]) were considered significantly varying and were excluded from the signature of bortezomib resistance. Note that some features above the red line in (E) vary between only one pair of wild-type clones and were therefore not necessarily excluded from the final signature.

**Figure 2-Supplement 4. Select features contributed to the signature of bortezomib resistance**

| High in resistant cells (14) | Low in resistant cells (31) |
|---|---|
| Cells_Correlation_K_DNA_AGP | Cells_AreaShape_Zernike_4_2 |
| Cells_RadialDistribution_ RadialCV_ER_2of4 | Cells_Correlation_Manders_Mito_ER |
| Cells_RadialDistribution_ RadialCV_Mito_ 1of4 | Cells_Correlation_Manders_Mito_RNA |
| Cells_RadialDistribution_ RadialCV_RNA_ 2of4 | Cells_Intensity_IntegratedIntensity_DNA |
| Cytoplasm_Correlation_K_DNA_AGP | Cells_Texture_Correlation__Mito_10_00 |
| Cytoplasm_Granularity_2_AGP | Cells_Texture_Correlation_Mito_10_02 |
| Cytoplasm_RadialDistribution_ MeanFrac_ Mito_3of4 | Cells_Texture_InfoMeas2_DNA_5_02 |
| Nuclei_AreaShape_Zernicke_6_0 | Cytoplasm_Correlation_K_AGP_DNA |
| Nuclei_Granularity_7_DNA | Cytoplasm_Correlation_Manders_ER_AGP |
| Nuclei_RadialDistribution_ MeanFrac_AGP_ 1of4 | Cytoplasm_Correlation_Manders_ER_RNA |
| Nuclei_RadialDistribution_ MeanFrac_AGP_ 2of4 | Cytoplasm_Correlation_Manders_RNA_AGP |
| Nuclei_RadialDistribution_ MeanFrac_RNA_ 1of4 | Cytoplasm_Correlation_RWC_DNA_ER |
| Nuclei_RadialDistribution_ RadialCV_ER_ 1of4 | Cytoplasm_Correlation_RWC_DNA_RNA |
| Nuclei_RadialDistribution_ RadialCV_RNA_ 2of4 | Cytoplasm_Correlation_RWC_Mito_ER |
| | Cytoplasm_Intensity_IntegratedIntensityEdge_RNA |
| | Cytoplasm_Intensity_MassDisplacement_ER |
| | Cytoplasm_Intensity_MassDisplacement_Mito |
| | Cytoplasm_Texture_AngularSecondMoment_Mito_5_02 |
| | Cytoplasm_Texture_InfoMeas2_Mito_5_00 |
| | Cytoplasm_Texture_InfoMeas2_Mito_5_01 |
| | Nuclei_AreaShape_Zernike_9_3 |
| | Nuclei_Texture_Correlation_DNA_10_00 |
| | Nuclei_Texture_Correlation_DNA_10_02 |
| | Nuclei_Texture_Correlation_DNA_5_00 |
| | Nuclei_Texture_Correlation_Mito_10_00 |
| | Nuclei_Texture_Correlation_Mito_10_01 |
| | Nuclei_Texture_Correlation_Mito_10_02 |
| | Nuclei_Texture_Correlation_RNA_10_01 |
| | Nuclei_Texture_Correlation_RNA_10_03 |
| | Nuclei_Texture_Corrleation_Mito_10_03 |
| | Nuclei_Texture_InfoMeas1_RNA_5_00 |

**Figure 2-Supplement 4. Select features contributed to the signature of bortezomib resistance.** List of 45 features that, after the exclusion of technical variables, were found to contribute to the signature of bortezomib resistance. Feature details can be found in the documentation for CellProfiler (see methods).

**Figure 2-Supplement 5. Bortezomib Signature-contributing features did not universally correlate**

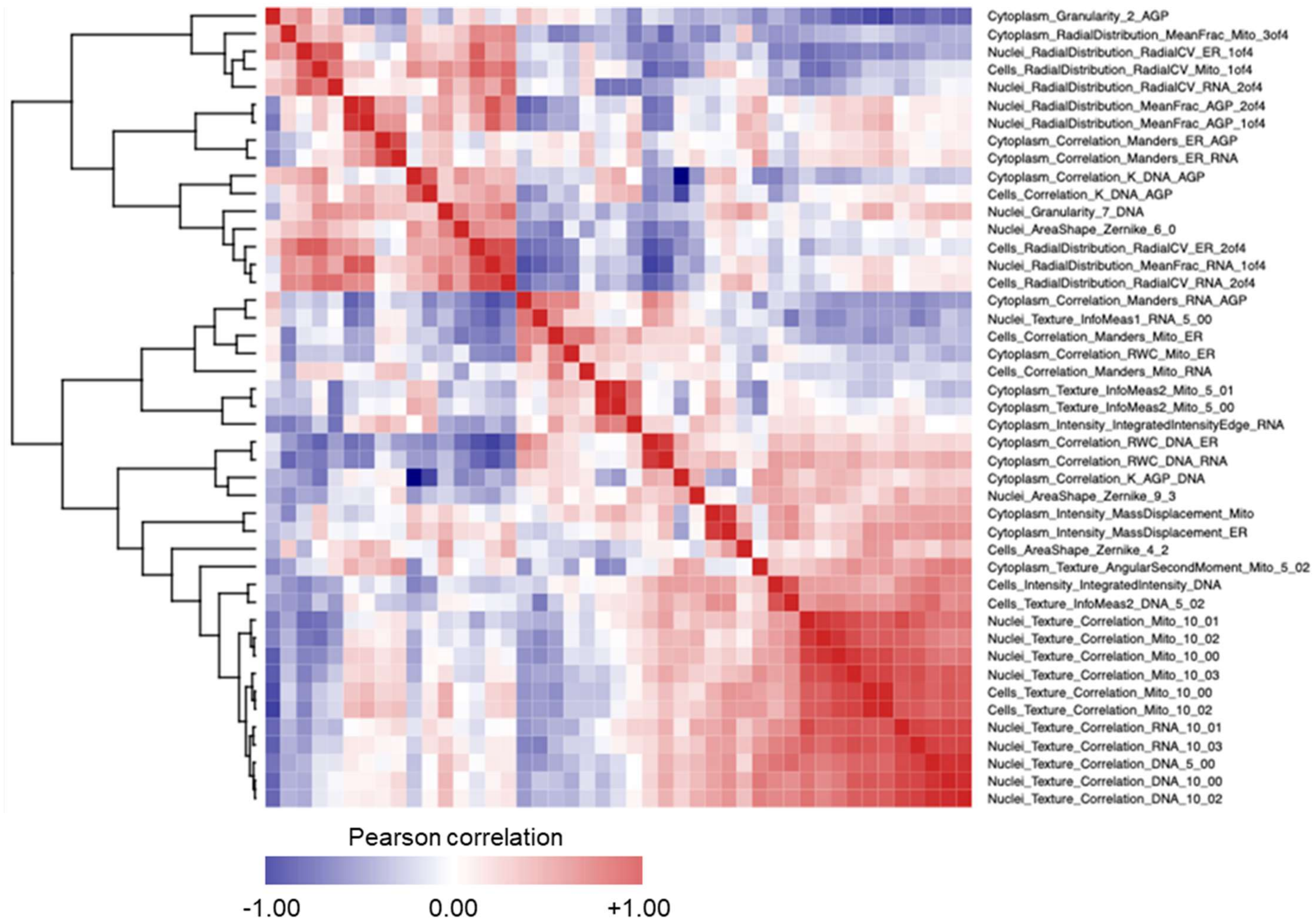


**Figure 2-Supplement 5. Bortezomib Signature-contributing features did not universally correlate.** Similarity matrix of pair-wise Pearson correlation coefficients of all 45 Bortezomib Signature features. Dendrogram displays hierarchical clustering of pairwise similarity.

**Figure 2-Supplement 6. Plating pattern does not strongly correlate with Bortezomib Signature**

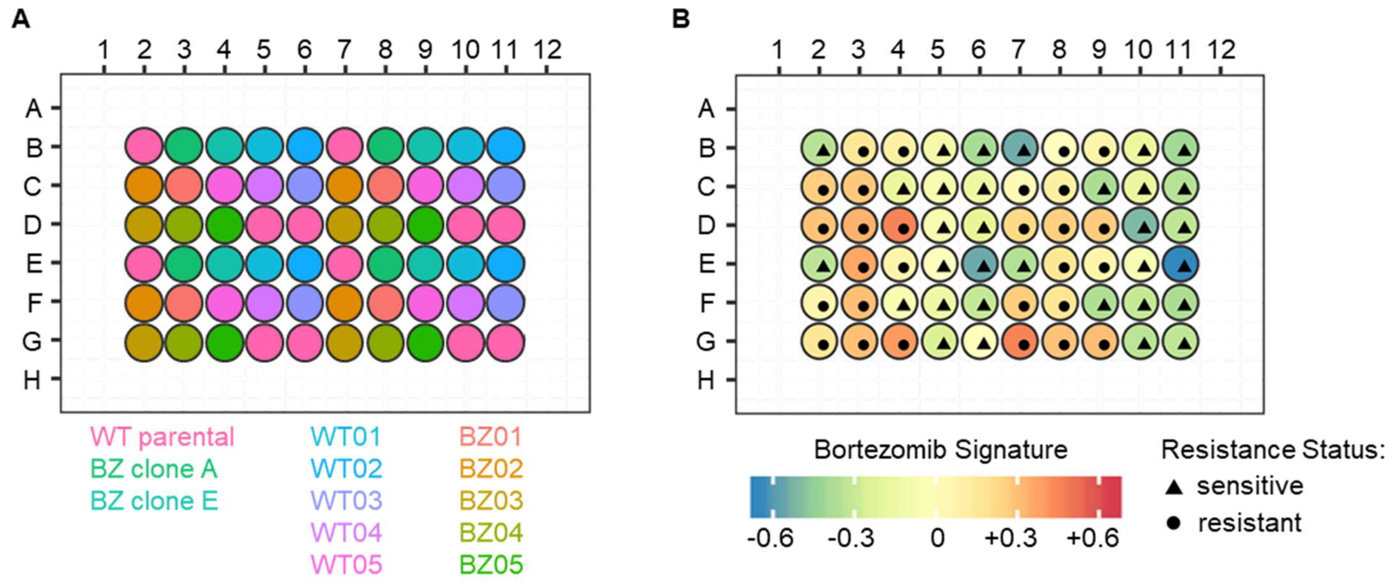


**Figure 2-Supplement 6. Plating pattern does not strongly correlate with Bortezomib Signature. (A)** Schematic of the plating pattern for HCT116 cell lines in a Cell Painting assay. Different cell lines are distinguished by color. **(B)** Bortezomib Signature for each cell line on the representative plate.

**Figure 2-Supplement 7. Bortezomib Signature identifies resistant clones and not technical variables**

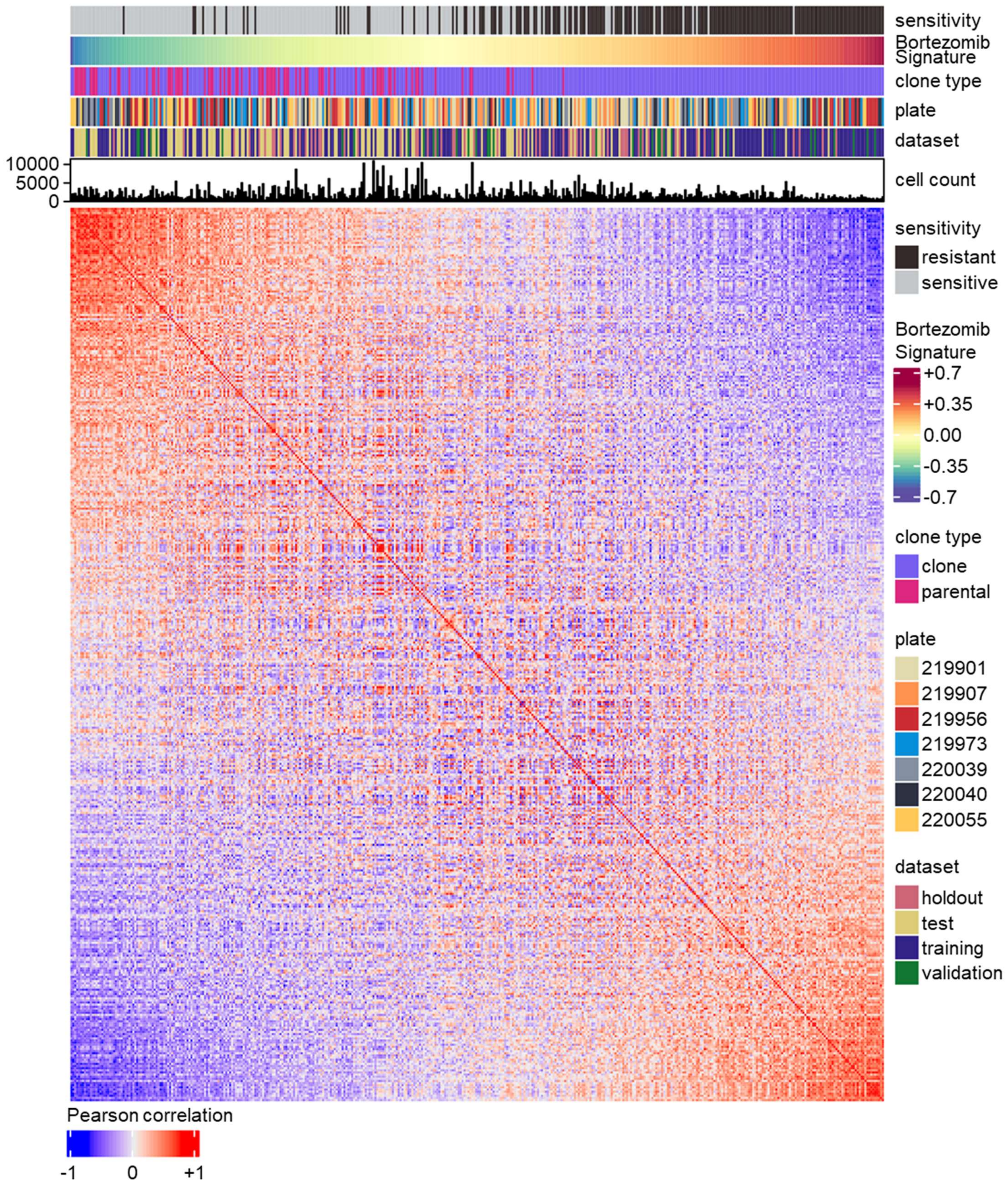

**Figure 2-Supplement 7. Bortezomib Signature identifies resistant clones and not technical variables.** Similarity matrix of Pearson correlation coefficients for cell line well-level profiles. Drug sensitivity, Bortezomib Signature, clone type (polyclonal parental or clonal), plate number (batch), dataset (for signature evaluation and generation), and cell count are indicated.

**Figure 2-Supplement 8. Datasets for Bortezomib Signature generation and evaluation**

| Clone ID | Training | Validation | Test | Holdout |
|---|---|---|---|---|
| WT parental | 0 | 0 | 72 | 12 |
| WT01 | 21 | 3 | 0 | 4 |
| WT02 | 20 | 4 | 0 | 4 |
| WT03 | 20 | 4 | 0 | 4 |
| WT04 | 20 | 4 | 0 | 4 |
| WT05 | 21 | 3 | 0 | 4 |
| BZ clone A | 0 | 0 | 24 | 4 |
| BZ clone E | 0 | 0 | 24 | 4 |
| BZ01 | 20 | 4 | 0 | 4 |
| BZ02 | 21 | 3 | 0 | 4 |
| BZ03 | 20 | 4 | 0 | 4 |
| BZ04 | 20 | 4 | 0 | 4 |
| BZ05 | 21 | 3 | 0 | 4 |

**Figure 2-Supplement 8. Datasets for Bortezomib Signature generation and initial evaluation.** Breakdown of the number of well-based profiles from cell lines used for the training, validation, test, and holdout datasets used to generate and evaluate the Bortezomib Signature. Data from 6 independent experiments contributed to the training, validation, and test datasets. The holdout dataset is one separate experiment. Note: signature evaluated on clones not included in the training dataset (WT10, WT12-WT15, BZ06-BZ10) in a separate analysis, see Figure 3.

**Figure 2-Supplement 9. Bortezomib Signatures from individual cell lines in training, validation, test, and holdout datasets**

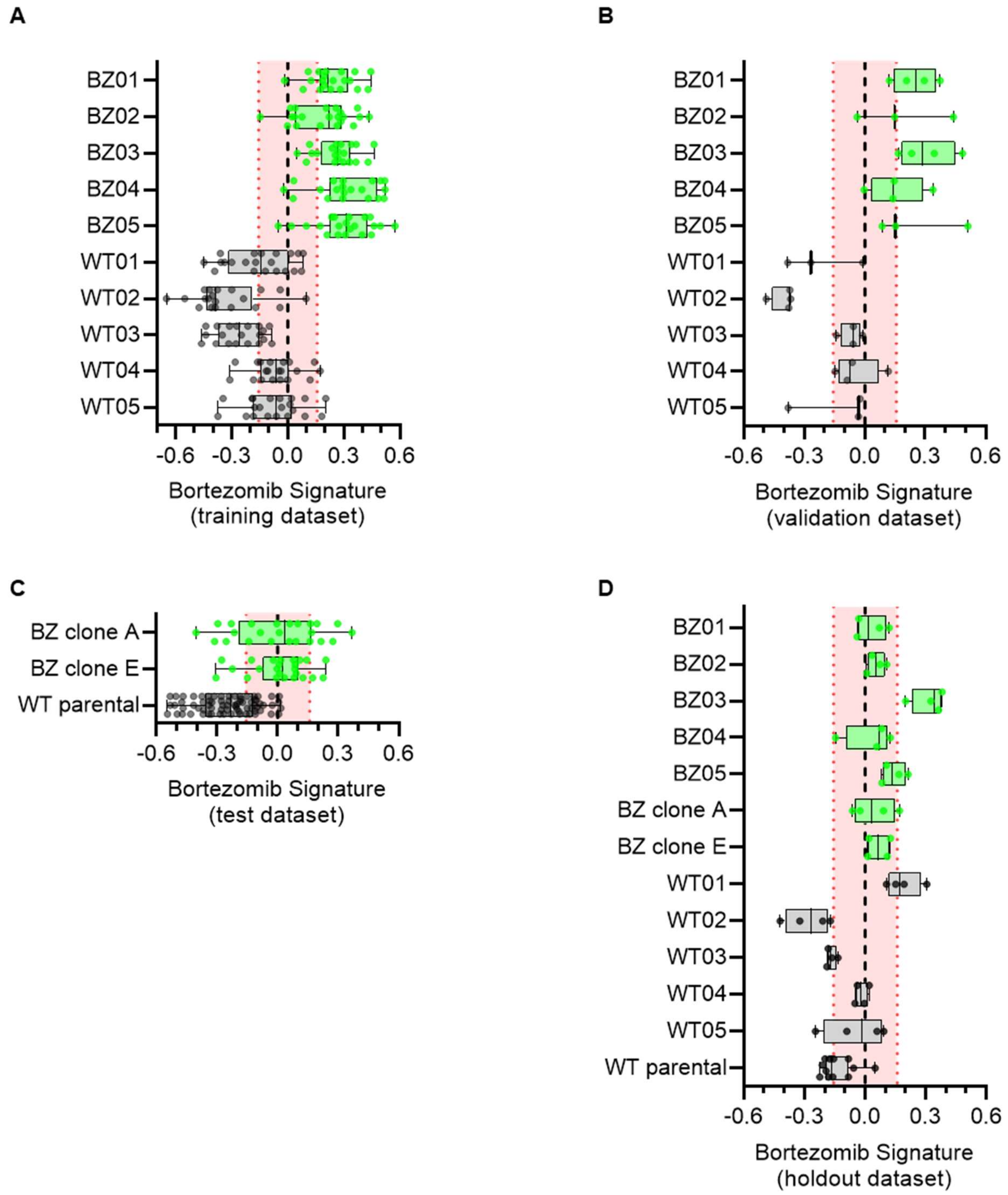


**Figure 2-Supplement 9. Bortezomib Signatures from individual cell lines in training, validation, test, and holdout datasets.** Box plots of Bortezomib Signatures for cell lines in the **(A)** training, **(B)** validation, **(C)** test, and **(D)** holdout datasets. Plots show individual points, range (error bars), 25th and 75th percentiles (box boundaries), and median. Dashed vertical black line is Bortezomib Signature = 0, dashed vertical red lines are the 95% confidence interval for Bortezomib Signatures of 1000 random permutations of the data. See Fig. 2-Supplement 8 for breakdown of profiles and experiments per dataset.
